# Innate extracellular Hsp70 inflammatory properties are mediated by the interaction of Siglec-E and LOX-1 receptors

**DOI:** 10.1101/2023.12.01.569623

**Authors:** Thiago J. Borges, Karina Lima, Ayesha Murshid, Isadora T. Lape, Yunlong Zhao, Maurício M. Rigo, Benjamin J. Lang, Shoib S. Siddiqui, Enfu Hui, Leonardo V. Riella, Cristina Bonorino, Stuart K Calderwood

**Author notes:** **Corresponding authors’ information:** Dr. Stuart K. Calderwood, BIDMC, 330 Brookline Ave, East Campus DA-717A, Boston, MA. 02215. Dr. Thiago J. Borges, Massachusetts General Hospital – Charlestown Navy Yard, 149 13^th^ St, room 5101A, Boston, MA. 02129.

## Abstract

Innate immune responses to cell damage-associated molecular patterns induce a controlled degree of inflammation, ideally avoiding the promotion of intense unwanted inflammatory adverse events. When released by damaged cells, Hsp70 can stimulate different responses that range from immune activation to immune suppression. The effects of Hsp70 are mediated through innate receptors expressed primarily by myeloid cells, such as dendritic cells (DCs). The regulatory innate receptors that bind to extracellular mouse Hsp70 (mHsp70) are not fully characterized, and neither are their potential interactions with activating innate receptors. Here, we describe that extracellular mHsp70 interacts with a receptor complex formed by inhibitory Siglec-E and activating LOX-1 on DCs. We also find that this interaction takes place within lipid microdomains, and Siglec-E acts as a negative regulator of LOX-1-mediated innate activation upon mHsp70 or oxidized LDL binding. Thus, HSP70 can both bind to and modulate the interaction of inhibitory and activating innate receptors on the cell surface. These findings add another dimension of regulatory mechanism to how self-molecules contribute to dampening of exacerbated inflammatory responses.

## Introduction

A key question in innate immunity is how cells distinguish between self- and foreign molecules with high degrees of homology. Innate immune responses are initiated by pattern-recognition receptors (PRRs), expressed in abundance by macrophages and dendritic cells (DCs). Their ligands vary from pathogen-associated molecular patterns (PAMPs) present in microorganisms to damage-associated molecular patterns (DAMPs) derived from injured tissues. Ideally, an effector immune response must be able to resolve any infection or sterile trauma with no exacerbating deleterious immunopathology. Importantly, the interplay of innate signals will shape the adaptive immune responses that ensue ^1^.

Heat shock protein (HSP) HSP70 is one of the most conserved evolutionarily proteins but has complex effects when present in the extracellular microenvironment. Mouse HSP70 (mHSP70) binds to scavenger receptors like LOX-1, SREC-I and FEEL-1, and triggers inflammatory innate immunity on DCs ^2^. DCs are the major antigen-presenting cells (APCs) in mammals and play a crucial role in inducing and regulating T cell responses, which initiate upon engaging MHC:peptide complexes and co-stimulatory molecules (CD86 and CD80) expressed by the APCs. Conversely, HSP70 can exhibit anti-inflammatory properties through its modulation of APCs and the induction of regulatory T cells (Tregs) ^3–7^. Consequently, extracellular HSP can either enhance or suppress immunity depending on the microenvironmental context. Recent findings demonstrated that human HSP70 could interact with paired receptors Siglec-5 and Siglec-14, with Siglec-14 enhancing IL-8 and TNF-α production in human monocytes following stimulation by human HSP70, while Siglec-5 reduced this inflammatory signaling ^8^. However, the mechanism by which HSPs could induce opposing responses in innate immune cells is poorly understood at a molecular level. One possible scenario is that the same HSP70 molecule could be recognized by different functional receptors to activate distinct innate responses. Alternatively, they could be recognized by the same receptor but lead to differential immune responses due to interaction with distinct co-receptors.

Here, we show that extracellular mHSP70 binds to mouse Siglec-E, a receptor recognized for its capacity to restrain inflammatory signaling. Notably, Siglec-E could also interact with the activating mHsp70 receptor LOX-1, forming a complex within lipid microdomains in the plasma membranes of DCs. Deletion of Siglec-E from DCs increased LOX-1-triggered activation, while its overexpression reduced LOX-1-mediated effects, suggesting that Siglec-E restricts LOX-1 engagement on DCs. Our results suggest a mechanism by which innate counter-receptors heterodimerize on the cell surface, forming a receptor complex that binds to non-glycosylated self-molecules, thus contributing to the intricate orchestration of immune responses.

## Results

### Extracellular mHSP70 interacts with Siglec-E

We first analyzed the potential role of Siglec-E – one of the main murine Siglec receptors - in mHSP70-mediated effects. We asked whether mHSP70 could bind directly to Siglec-E. As a first approach, we generated a murine Siglec-E receptor construct encoding an HA tag in its N-terminal (extracellular) domain. We then transfected CHO-K1 cells with Siglec-E-HA plasmids and analyzed the binding and co-localization of the expressed receptor with fluorescently tagged mHSP70 in the absence of other potential receptors. Notably, wild-type CHO-K1 cells do not bind extracellular Hsp70 based on previous screening studies, providing a clean platform to examine the abilities of innate receptors to bind and respond to HSP70 ^9,10^. Alexa 488-tagged mHSP70 bound to CHO-K1 expressing Siglec-E, but not to the untransfected cells (**Fig. 1A**). Such binding was inhibited by an anti-Siglec-E antibody (**Fig. 1A**). By confocal microscopy, we confirmed both that mHSP70 did not bind to CHO-K1 control cells, and that it localized with HA (red, merged represented by yellow area) in Siglec-E-HA-expressing CHO cells (**Fig 1B**). To further confirm these findings, ELISA plates were coated with purified mHSP70 and probed with a recombinant soluble IgG-Fc fusion chimera containing the extracellular domain of Siglec-E (**Fig 1C**). mHSP70 interacted directly with Siglec-E under these conditions (**Fig 1D**) confirming the binding of the HSP70 to Siglec-E. We then examined HSP70 binding in primary DCs *ex vivo*. We incubated DCs isolated from naïve mice spleens with extracellular Alexa 488-tagged mHSP70 and evaluated its potential co-localization with Siglec-E on the cells. mHSP70 co-localized with Siglec-E in the murine DCs as indicated by areas of the yellow merged fluorescence (**Fig 1E**), with a Pearson’s correlation of 0.702 ± 0.127. In conclusion, extracellular mHSP70 can bind to murine Siglec-E.

**Fig 1.**
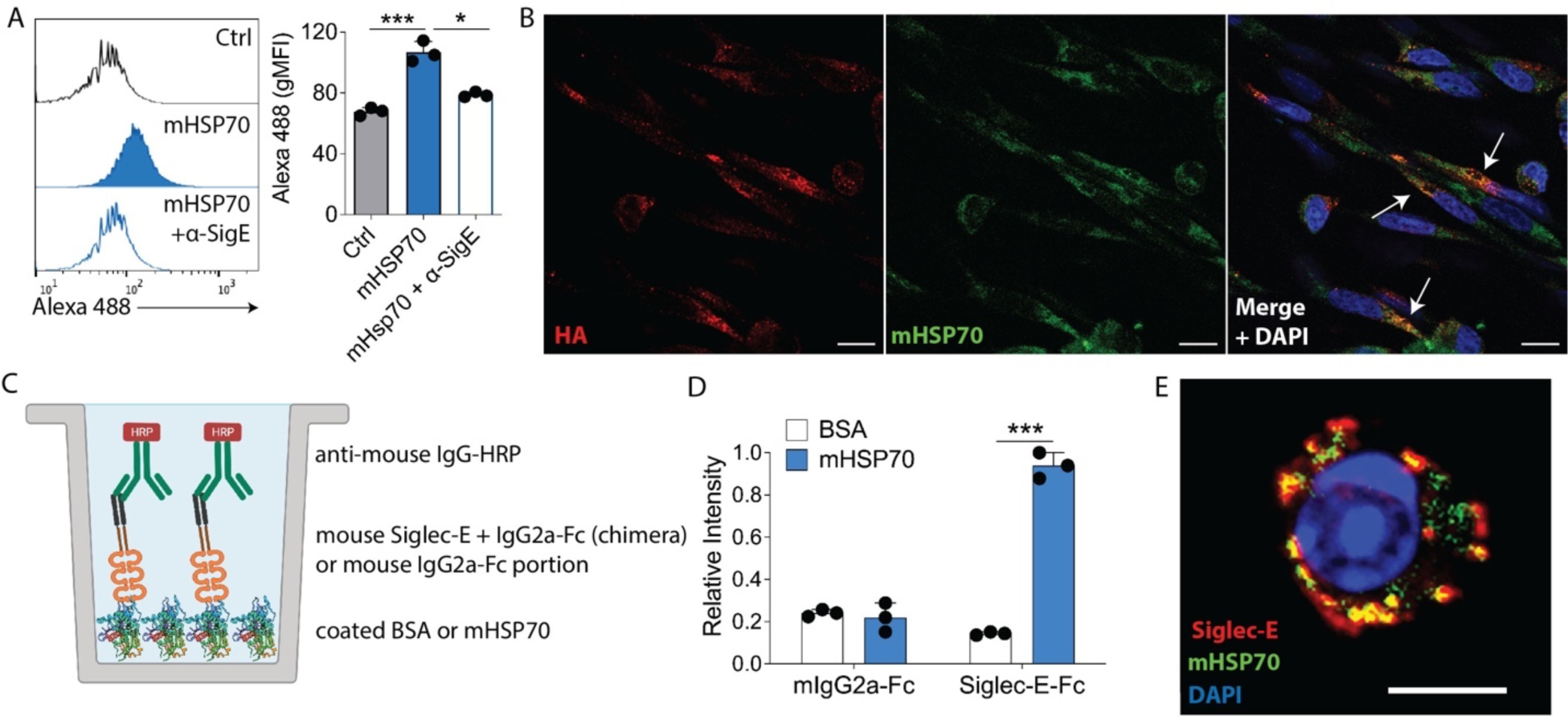
mHSP70 binds to murine Siglec-E. (**A**) CHO cells stably overexpressing Siglec-E-HA were treated with Alexa 488-tagged murine HSP70 (mHSP70) on ice for 30 min and its binding was evaluated by flow cytometry. Anti-Siglec-E blocking antibody was added to neutralize the binding. *p<0.05 or ***p<0.001 when compared to mouse IgG using one-way ANOVA with Tukey post-hoc test. gMFI: geometric mean fluorescence intensity. (**B**) Representative confocal microscopy images of CHO cells stably overexpressing Siglec-E-HA and treated with Alexa 488-tagged mHSP70. Cells were then stained for HA (red). Magnification 400x. Scale bar = 5 μm (**C**) Representation of the ELISA in which plates were coated with mHSP70 or BSA (negative control), followed by incubation with a Siglec-E-Fc chimera and a secondary HRP anti-mouse IgG. Representation created with BioRender.com. (**D**) Relative binding between mHSP70 to Siglec-E-Fc determined by ELISA. ***p<0.001 when compared to BSA by t-test. (**E**) Representative confocal image of mHSP70 (green) binding to Siglec-E (red) in splenic DCs isolated from naïve WT mice with magnetic CD11c beads. Scale bar = 10 μm. Bars are represented as mean ± SD of triplicates. All data are representative of at least three independent experiments.

### mHSP70 is a non-sialylated ligand for Siglec-E

Siglec-E is known to bind specifically to glycosylated proteins. We thus asked whether the mHSP70 used in the study was glycosylated. Our mHSP70 preparation did not contain detectable levels of N-glycans (**Fig 2A**), when compared with the heavily glycosylated fetuin, used as a positive control (**Fig 2B**). We confirmed these findings by SDS-PAGE and Western Blot analyses. Deglycosylation of mHSP70 with the N-glycosidase PNGase F treatment did not result in a band shift in SDS-PAGE analysis, contrary to what was observed with fetuin (**Fig 2C**). Additionally, the presence of putative α(2,6) sialoglycans displayed on mHSP70 was further analyzed. Western blots showed that the lectin SNA was not able to bind to mHSP70 (**Fig 2D**). These findings led us to hypothesize that Siglec-E recognizes an alternative peptide-based topology on the HSP molecule, unrelated to glycosylation.

**Fig 2.**
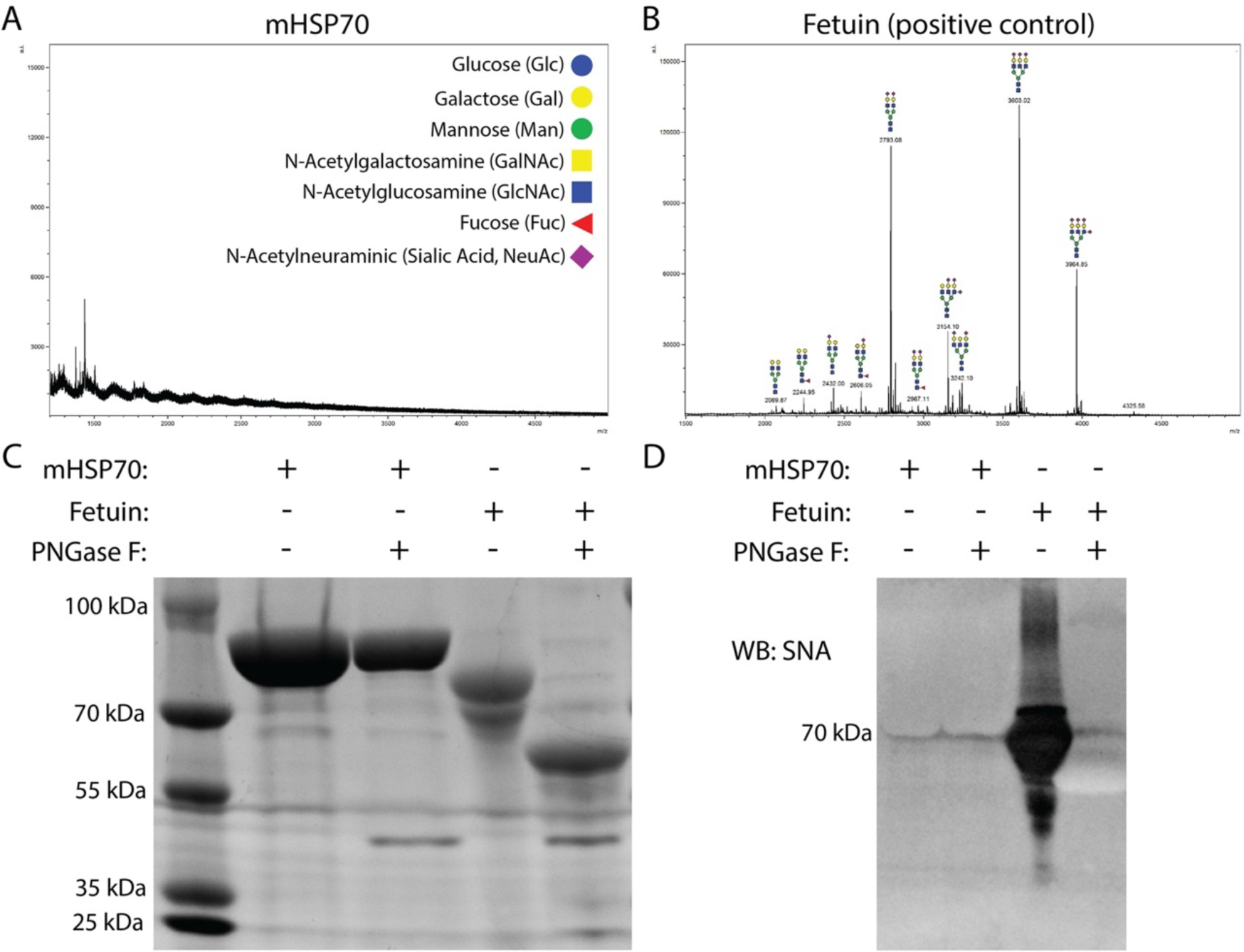
Mouse Hsp70 is not glycosylated. (**A**) Analysis of N-glycans profile of extracellular mHsp70. (**B**) Fetuin was used as a positive control. MALDI-TOF mass spectrometry was performed in protein preparations in which the N-linked glycans were released enzymatically by PNGase F and permethylated, before profiled. (**C**) SDS-PAGE of mHSP70 or fetuin treated or not with PNGase F (deglycosylation). (**D**) SNA lectin blot analysis of purified extracellular mHSP70 or fetuin (positive control) treated or not with PNGase F. (**C** - **D**) One representative gel from n = 2 biologically independent experiments.

### Siglec-E and LOX-1 form innate receptor complexes in immune cells

Mammalian HSP70 has been previously demonstrated to bind to the innate receptor LOX-1 ^9,11^. We asked if Siglec-E and LOX-1 could interact and potentially form complexes in DCs, cooperating to fine-tune opposing signals, and consequently their inflammatory/anti-inflammatory effects. To test this hypothesis, we coated plates with purified LOX-1 and probed them with different concentrations of a soluble Siglec-E-Fc chimera (**Fig 3A**). An ELISA revealed that the binding occurred in a dose-dependent manner (**Fig 3B**) and was partially inhibited by an anti-LOX-1 antibody (**Fig 3C**). We then confirmed their interaction by immunoprecipitating Siglec-E from lysed murine splenocytes. Using immunoblot analysis, we detected considerable amounts of LOX-1 in the Siglec-E immunoprecipitate (**Fig 3D**). Interestingly, Siglec-E is also associated with endogenous mHSP70 present in the splenocytes (**Fig 3D**). We confirmed these findings by transfecting stable CHO-LOX-1-Myc cells with Siglec-E-HA plasmids and analyzing their interaction by confocal microscopy. When we transfected Siglec-E into CHO cells expressing LOX-1, we noticed that surface expression patterns changed dramatically when compared to wild-type CHO transfectants (**Fig 3E**). In CHO cells expressing only Siglec-E, receptor distribution seemed more evenly spread over the cell surface. However, Siglec-E expression became concentrated into foci in cells when co-transfected with LOX-1 (**Fig 3E**), suggesting an association in discrete patches on the membrane. By staining for HA and Myc tags, we observed that Siglec-E and LOX-1 colocalization could be detected in discrete foci in the CHO cells (arrows, **Fig 3F**). To further characterize Siglec-E/LOX-1 interaction in intact cells, we examined their co-localization in non-stimulated DCs isolated from murine spleens. Indeed, flow cytometry showed that ∼15% of CD11c^hi^ cells co-expressed Siglec-E and LOX-1 on their cell surface, but not in Siglec-E KO cells as expected (**Fig 3G**). Overall, these results support a significant interaction between Siglec-E and LOX-1 on DCs.

**Fig 3.**
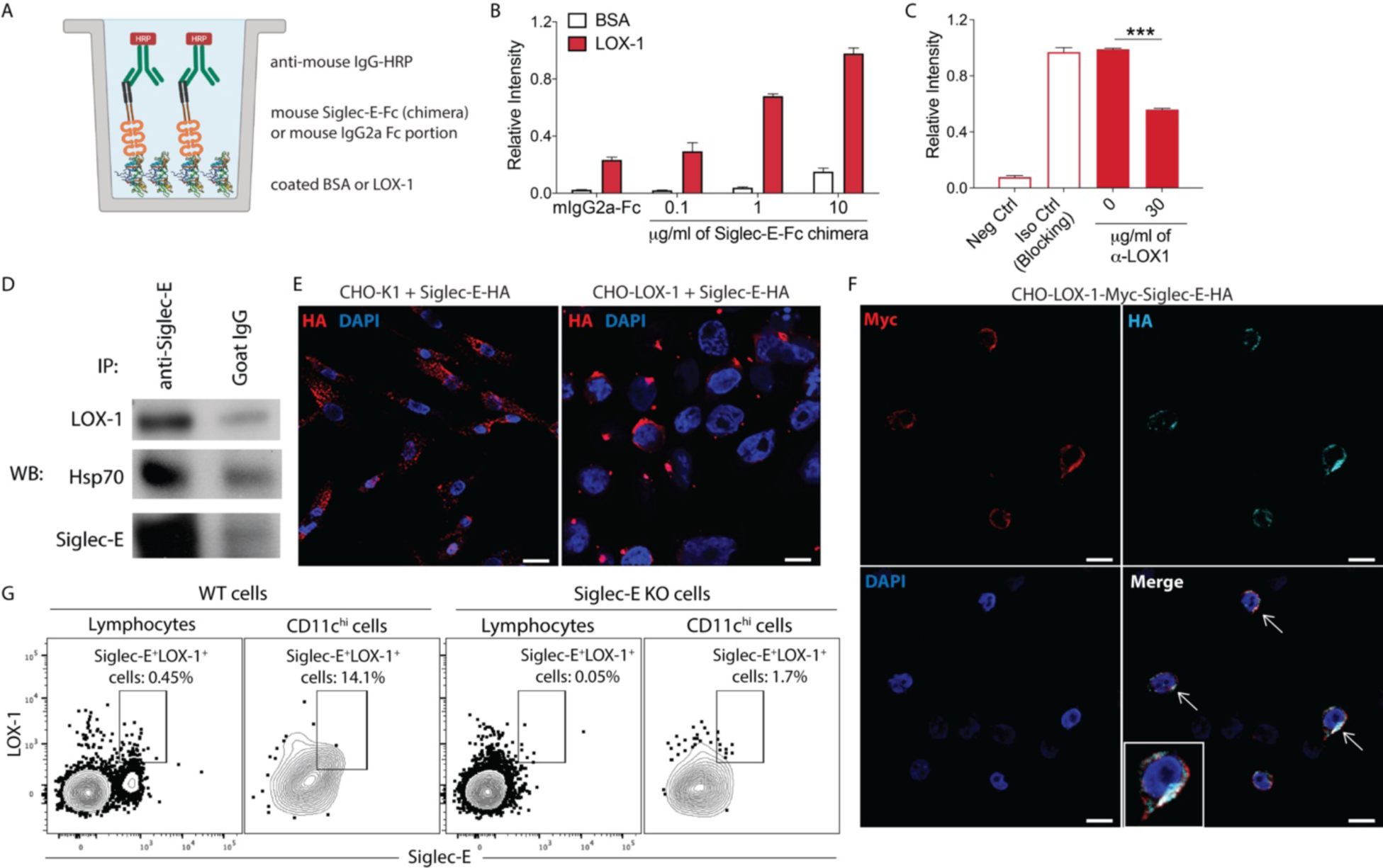
Innate receptors Siglec-E and LOX-1 form hetero-complexes and are co-expressed by murine CD11c^hi^ DCs. (**A**) Cartoon of the ELISA in which LOX-1 or BSA (negative control) were coated in the plate, incubated initially with different concentrations of a soluble Siglec-E-Fc chimera, followed by incubation with a secondary HRP anti-mouse IgG. Representation created with BioRender.com. (**B**) Relative binding between LOX-1 and Siglec-E-Fc chimera is dose-dependent. All assays were performed in triplicates. (**C**) Relative binding between LOX-1 and Siglec-E-Fc chimera in the presence of anti-LOX-1 blocking antibody or isotype control. ***p<0.001 by one-way ANOVA with Tukey post-test. All assays were performed in triplicates. (**D**) Lysates from WT naïve spleen cells were precipitated with either Siglec-E–specific antibody or control mouse IgG. Precipitates were probed with antibodies to Siglec-E, LOX-1 or mHSP70. Complete membrane blots are included in Supplementary Fig. 1. One representative gel from n = 2 biologically independent experiments. (**E**) CHO-K1 cells or CHO cells stably overexpressing LOX-1 were transfected with a plasmid containing the Siglec-E-HA construct. Siglec-E expression pattern (HA staining, red) was analyzed by confocal microscopy. (**F**) Representative confocal microscopy images of untreated CHO cells stably overexpressing LOX-1-Myc and Siglec-E-HA receptors stained for c-Myc (red) and HA (cyan). (**E** – **F**) Magnification 400x. Scale bar = 5 μm. All data are representative of three independent experiments. (**G**) Representative contour plots of Siglec-E^+^LOX-1^+^ cells in total lymphocytes or CD11c^high^ (DCs) cells from WT or Siglec-E KO naïve mice spleens. Representative plots from n = 3 biologically independent experiments.

### Molecular docking and electrostatic potential analysis of Siglec-E and LOX-1 interaction

We next employed complementary bioinformatic analysis to further explore the molecular nature of the interactions between Siglec-E and LOX-1. We first generated *in silico* 3D models for both Siglec-E and LOX-1 receptors. The two best templates for Siglec-E were a myelin-associated glycoprotein (MAG) from *Mus musculus* (PDB ID: 5LFR) and two N-terminal domains of SIGLEC-5 from *Homo sapiens* (PDB ID: 2ZG2). The whole extracellular moiety comprising the residues 19 to 353 of Siglec-E was modeled using the multiple templates approach from *Modeller*. Regarding the murine LOX-1 protein (target), the best template structure found was the LOX-1 molecule from *Homo sapiens* (PDB ID: 1YPQ). Previous data indicate that mouse LOX-1 possesses the structural features required to form dimers, as it has been shown to occur with homologous human LOX-1 ^12^. Also, the C-terminal C-type lectin-like domain (CTLD) is highly conserved among species and was already shown to be involved in the binding of several molecules by C-type lectins such as LOX-1 ^13,14^. For this reason, we decided to model the CTLD from LOX-1 as a homodimer and analyzed *in silico* its interaction with Siglec-E. Data from *ClusPro*, a well-known molecular docking server used to predict protein-protein interaction ^15^, predicted that a portion of the Ig-like V-domain from Siglec-E could interact with the CTLD from LOX-1 dimers (**Fig 4A-B**). Of potential significance, there is a charge complementarity at the interaction surface between both molecules, where two positively charged motifs in Siglec-E interact with a negatively charged region of CTLD from LOX-1 (**Fig 4C-D**). We screened residues present at the Siglec-E/LOX-1 dimer interface, focusing on those involved in hydrogen bonding (**Fig 4E**). Most of the predicted interacting Siglec-E residues are positively charged, including – ARG34, LYS32, ARG75, and ARG86 – while the LOX-1 residues are negatively charged – ASP112, GLU118, GLU255, and ASN256 (**Supplementary Table 1**). We identified additional residues that could be involved in the interaction (**Supplementary Table 1**). Hydrogen bonding between the molecules appeared to involve ARG34, ARG75 and ARG86 from Siglec-E and GLU112, GLU118, GLU255 and SER60 from LOX-1. In conclusion, our modeling studies suggest potential electrostatic interactions could occur between Siglec-E and the LOX-1 within their extracellular domains, along with additional hydrogen bonds, which would be crucial to maintaining a heterodimer.

**Fig 4.**
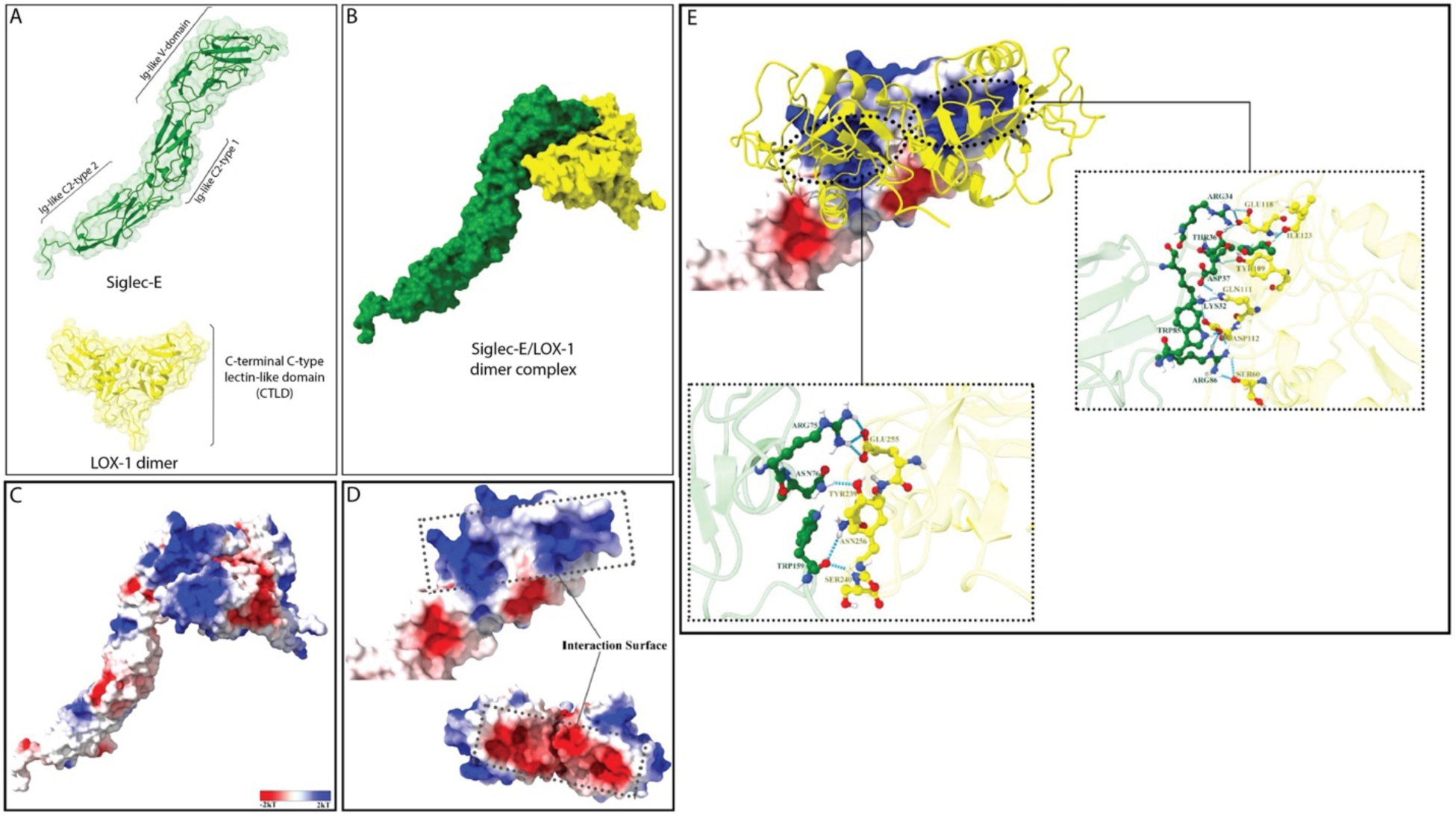
*In silico* modeling of Siglec-E and LOX-1 dimer and molecular docking and electrostatic potential analysis of the Siglec-E/LOX-1 dimer complex. Models of Siglec-E (green) and LOX-1 dimer (yellow) domains are represented in (**A**). In (**B**), a representation of Siglec-E and LOX-1 predicted interaction was obtained from the ClusPro web server. (**C**) Complex with electrostatic potential computed on the surface, where red, white and blue represent negative, neutral and positive charges, respectively (gradient ranges from -2kT to 2kT, being *k* the Boltzmann constant and *T* the temperature). (**D**) The molecules were separated and rotated to show the interaction surface between Siglec-E and LOX-1, evidencing a complementarity of charges. (**E**) LOX-1 is positioned exactly over the two positive charges of Siglec-E and the complex is stabilized by several hydrogen bonds (*cyan blue*) between Siglec-E (*green*) and LOX-1 (*yellow*).

### mHSP70 binds to Siglec-E/LOX-1 complexes present in lipid rafts

We next investigated whether mHSP70 could bind to the receptor complex formed by Siglec-E and LOX-1. To determine whether mHSP70 simultaneously binds to the receptor complex, we treated CHO-LOX-1-Myc-Siglec-E-HA cells with Alexa 488-tagged mHSP70. Fluorescently tagged mHSP70 co-localized with Siglec-E/LOX-1 complexes in CHO cells (**Fig 5A**). We confirmed these findings by treating splenic primary DCs with fluorescently tagged mHSP70 at 4°C to reduce endocytosis. mHSP70 bound to murine DCs and became co-localized with Siglec-E and LOX-1 (**Fig 5B**). Foci of co-localization (shown in white) were seen on the cell surface upon incubation with mHSP70, suggesting the presence of triple complexes between each mHsp70 (green), Siglec-E (cyan) and LOX-1 (red) (**Fig 5B**). We also observed that mHSP70, Siglec-E and LOX-1 complexes were internalized when conducting the binding assays at 37°C (**Fig 5C**). These complexes are found at a physiological level since the stain of endogenous mHSP70 (green) co-localized with Siglec-E (cyan) and LOX-1 (red) on the membrane of murine DCs (**Fig 5D**).

**Fig 5.**
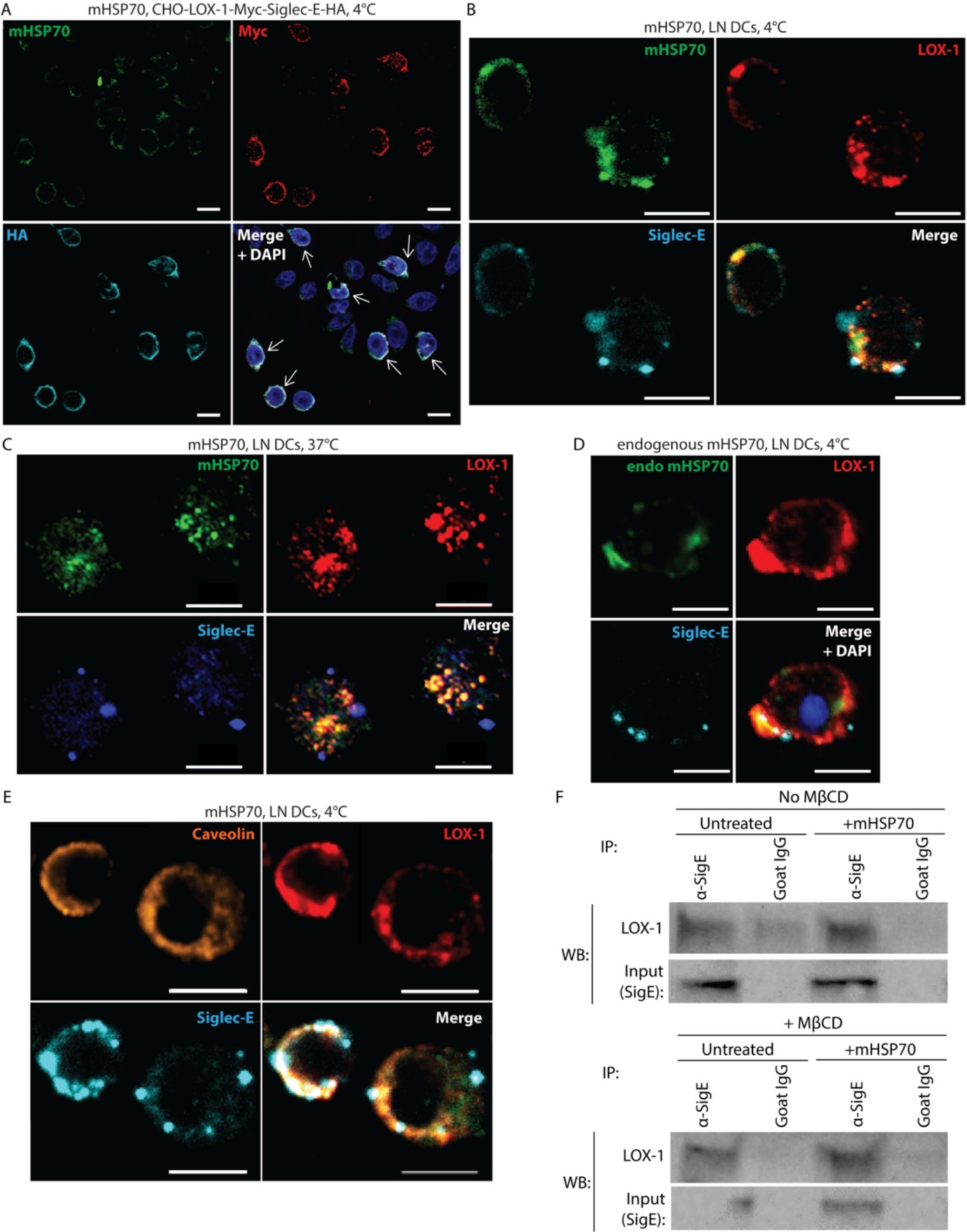
Siglec-E/LOX-1 complexes are localized in lipid microdomains. (**A**) CHO-LOX-1-Myc-Siglec-E-HA expressing cells were treated with Alexa 488-tagged extracellular murine HSP70 (mHSP70, 10 μg/ml) on ice for 30 min. Cells were then stained for c-Myc (red) and HA (Cyan) and analyzed by confocal microscopy. Representative confocal microscopy images of murine DCs isolated from naïve WT animals and treated with Alexa 488-tagged mHSP70 on ice (**B**) or at 37°C (**C**) for 30 min. Cells were then stained for LOX-1 (red) and Siglec-E (Cyan). (**D**) Representative confocal microscopy images of murine DCs stained for endogenous mHSP70 (green), LOX-1 (red) and Siglec-E (Cyan). (**E**) Representative confocal microscopy images of murine DCs stained for caveolin (orange), LOX-1 (red) and Siglec-E (Cyan) (**A** – **E**) Magnification 400x. Scale bars = 5 μm. (**F**) WT splenocytes were treated for 2h with MβCD (20 mM, cholesterol sequestering agent) before stimulation with mHSP70 for 30min. Spleen cell lysates were precipitated with either Siglec-E–specific antibody or control goat IgG. Precipitates were probed with antibodies specific to LOX-1 or Siglec-E (input). One representative gel from n = 3 biologically independent experiments. Complete membrane blots are included in Supplementary Fig. 1.

The focal concentrations of the Siglec-E/LOX-1 receptor complexes suggested that their association could occur in discrete patches on the membrane. Our previous studies on the Hsp70-binding scavenger receptors indicated that SRECI, a close functional homolog of LOX-1, was present in cholesterol-rich lipid rafts, and could influence other immune signaling molecules to enter these lipid microdomains and regulate their activities ^16^. Therefore, we investigated the possibility that LOX-1 might possess a similar property compared to its scavenger receptor co-member SREC-I. Indeed, Siglec-E (cyan) and LOX-1 (red) co-localized with the lipid raft marker Caveolin-1 (orange) in splenic mouse DCs (**Fig 5E**). We further investigated whether Siglec-E and LOX-1 interaction would increase upon stimulus with mHSP70 by calculating different colocalization coefficients. Compared to untreated cells, DCs stimulated with mHSP70 presented significantly higher co-localization between Siglec-E and LOX-1, as demonstrated by a higher Pearson’s correlation and Spearman’s rank correlation values (**Supplementary Table 2**).

To evaluate the potential role of lipid microdomains in stabilizing Siglec-E/LOX-1 complexes, we incubated WT splenocytes without or with cholesterol sequestering agent methyl β cyclodextrin (MβCD), known to disrupt lipid microdomains. We then treated the cells with mHSP70, and immunoprecipitated Siglec-E from the lysates of the treated cells, probing for the presence of LOX-1 by immunoblot. A slight increase in LOX-1 co-precipitate was observed in lysates treated with mHSP70, an effect that was partially blocked in cells pre-treated with MβCD (**Fig 5F**). These data supported the conclusion that Siglec-E/LOX-1 complexes are concentrated within lipid microdomains.

### Siglec-E controls the magnitude of the immune responses triggered by LOX-1 ligands

We next asked whether Siglec-E was involved in regulating the responses triggered by LOX-1 ligands. We tested the activation status of murine WT or Siglec-E-deficient or bone-marrow-derived DCs (BMDCs) incubated with increasing doses of mHSP70 or oxidized LDL (oxLDL) – a classic LOX-1 ligand ^17^. To exclude confounding effects of other contaminating molecular patterns, we analyzed the effects of endotoxin-free preparations of BMDCs by measuring MHC II, CD86, and CD80 levels, as well as the cytokines induced. Both oxLDL and mHSP70 increased the expression of MHC II (**Fig 6A**), CD86 (**Fig 6A**), and CD80 (**Fig 6C**) levels in a dose-dependent manner. This elevation was notably more significant in the absence of Siglec-E (**Fig 6A-C**). Analysis of unstimulated Siglec-E KO BMDCs indicated elevated basal levels of MHC II^hi^CD86^hi^ cells (**Fig 6D**), suggesting a different level of basal DC maturation state in these cells. This finding is consistent with the anti-inflammatory effects reported for Siglec-E ^18^. The increase in MHC II^hi^CD86^hi^ cells induced by oxLDL or mHSP70 was also more pronounced in Siglec-E KO BMDCs (**Fig 6D**). We next analyzed the production of cytokines in BMDCs supernatants using Luminex. The absence of Siglec-E significantly increased the production of IL-6 and TNF-α induced by oxLDL or mHSP70 (**Fig 6E**). Moreover, mHSP70 was able to induce the production of IL-18 by Siglec-E KO BMDCs, but not WT cells (**Fig 6E**). Interestingly, IL-10 production was higher in WT controls when compared to Siglec-E KO cells (**Fig 6E**). Thus, the absence of Siglec-E potentiates the expression of maturation markers and inflammatory cytokines production by BMDCs stimulated with LOX-1 ligands.

**Fig 6.**
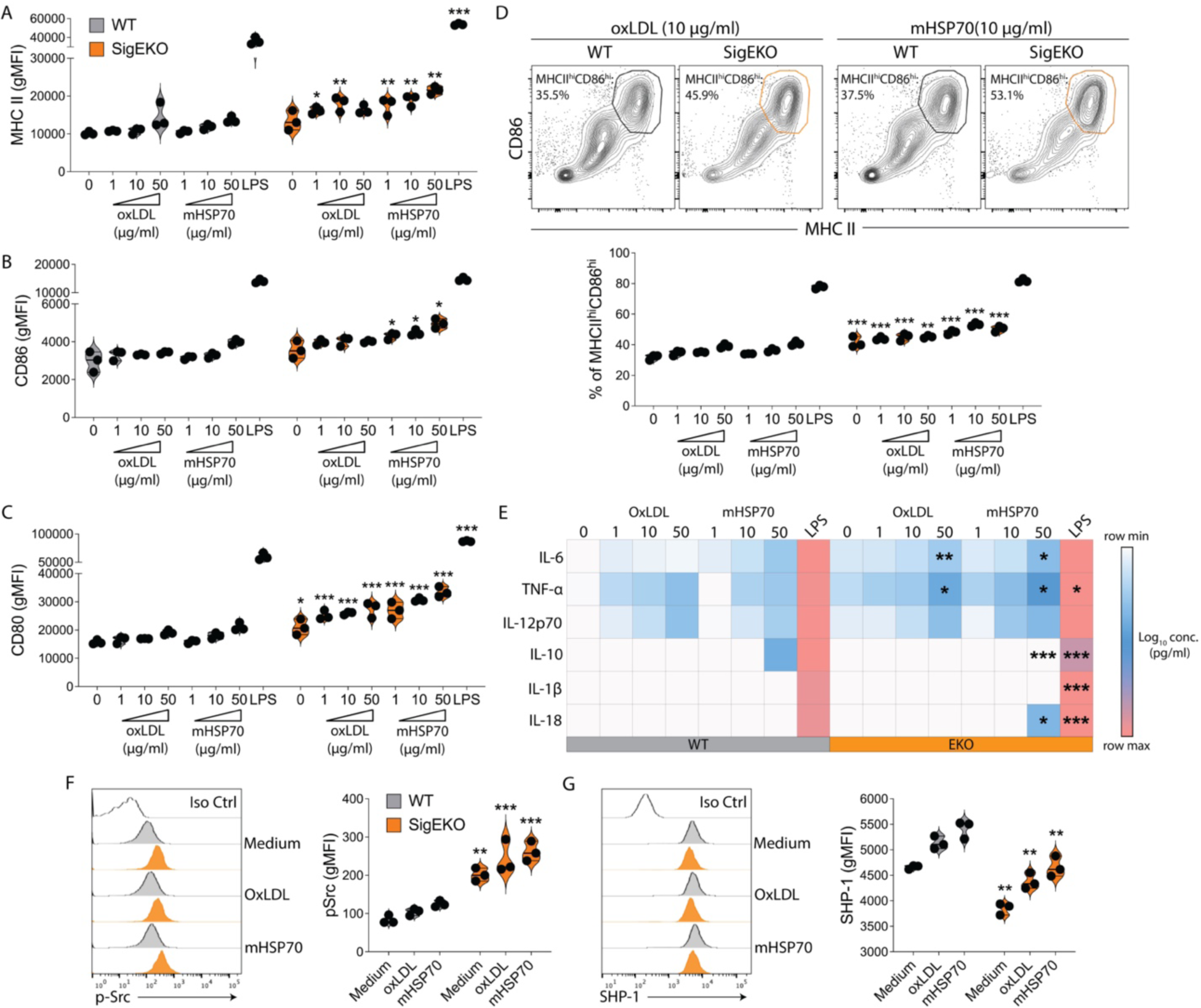
The absence of Siglec-E increases LOX-1-mediated activation of DCs. WT or Siglec-E KO BMDCs were treated with increasing doses of oxLDL or mHSP70 for 24h and (**A**) MHC II, (**B**) CD86, (**C**) CD80, or (**D**) MHC II^hi^CD86^hi^ cells were evaluated by flow cytometry. *p<0.05, **p<0.01, ***p<0.001 when compared to WT by one-way ANOVA with Tukey post-test. (**E**) Cytokine profile from culture supernatants analyzed by LUMINEX. The experiments are representative of two repetitions performed in triplicates of a pool of 2 mice per genotype. *p<0.05, **p<0.01, ***p<0.001 when compared to WT by one-way ANOVA with Tukey post-test. WT or Siglec-E KO BMDCs were treated with 50 μg/ml of oxLDL or 50 μg/ml of mHSP70 for 30 min and the expression of (**F**) phospho-Src (p-Src) or (**G**) SHP-1 was assessed by flow cytometry. **p<0.01, ***p<0.001 when compared to WT by one-way ANOVA with Tukey post-test. gMFI: geometric mean fluorescence intensity; Iso Ctrl: isotype control.

A prediction from these findings was that Siglec-E could potentially exert a negative regulatory effect on LOX-1-triggered intracellular signaling pathways. Given that LOX-1 signals through phosphorylated Src (p-Src) in DCs ^19^, we explored whether the impact on pSrc signaling is mediated by Siglec-E. In the absence of any ligand, Siglec-E KO BMDCs had increased baseline levels of p-Src compared to WT cells (**Fig 6F**). The treatment with either oxLDL or mHSP70 led to increased p-Src levels (**Fig 6F**). The upregulation of p-Src was notably more significant in Siglec-E-deficient DCs (**Fig 6F**). Siglec-E is known to recruit SH2-domain-containing protein tyrosine phosphatases SHP-1 and SHP-2 through its tyrosine-based inhibitory motifs (ITIM) motif and tyrosine-based switch motif (ITSM) ^20^, thus dampening inflammatory responses ^18,21^. Hence, we examined the levels of SHP-1 upon oxLDL or mHSP70 treatments. As expected, Siglec-E KO BMDCs have lower baseline levels of SHP-1 (**Fig 6G**). Although oxLDL and mHSP70 increased SHP-1 levels in Siglec-E-deficient DCs, this was significantly reduced when compared to WT cells (**Fig 6G**).

We next investigated whether the overexpression of Siglec-E on DCs would inhibit LOX-1-triggered DC activation. For this purpose, we utilized the DC cell line DC2.4, which express significant levels of LOX-1 (**Fig 7A**). We generated DC2.4 cells overexpressing Siglec-E (DC2.4 SigE^+^, **Fig 7B**). Both cell lines were treated with either LOX-1 ligand oxLDL, mHSP70 or LPS and DC activation was determined by evaluating the MHC II, CD86 and CD80 levels, as well as the cytokine profile by flow cytometry. We observed that WT DC2.4 cells had a significantly greater increase in the expression of MHC II (**Fig 7C**), CD86 (**Fig 7D**), and CD80 (**Fig 7E**) when compared to DC2.4 SigE^+^ cells. In addition, the increase in the production of TNF-α (**Fig 7F**) and IL-6 (**Fig 7G**) was lower in DC2.4 SigE^+^ cells when compared with the WT. Thus, our data suggest that overexpression of Siglec-E inhibits DC activation induced by both self (oxLDL, mHSP70) and foreign (LPS) ligands. Taken together, our data demonstrate that Siglec-E regulates, and fine-tunes inflammatory effects induced via LOX-1.

**Fig 7.**
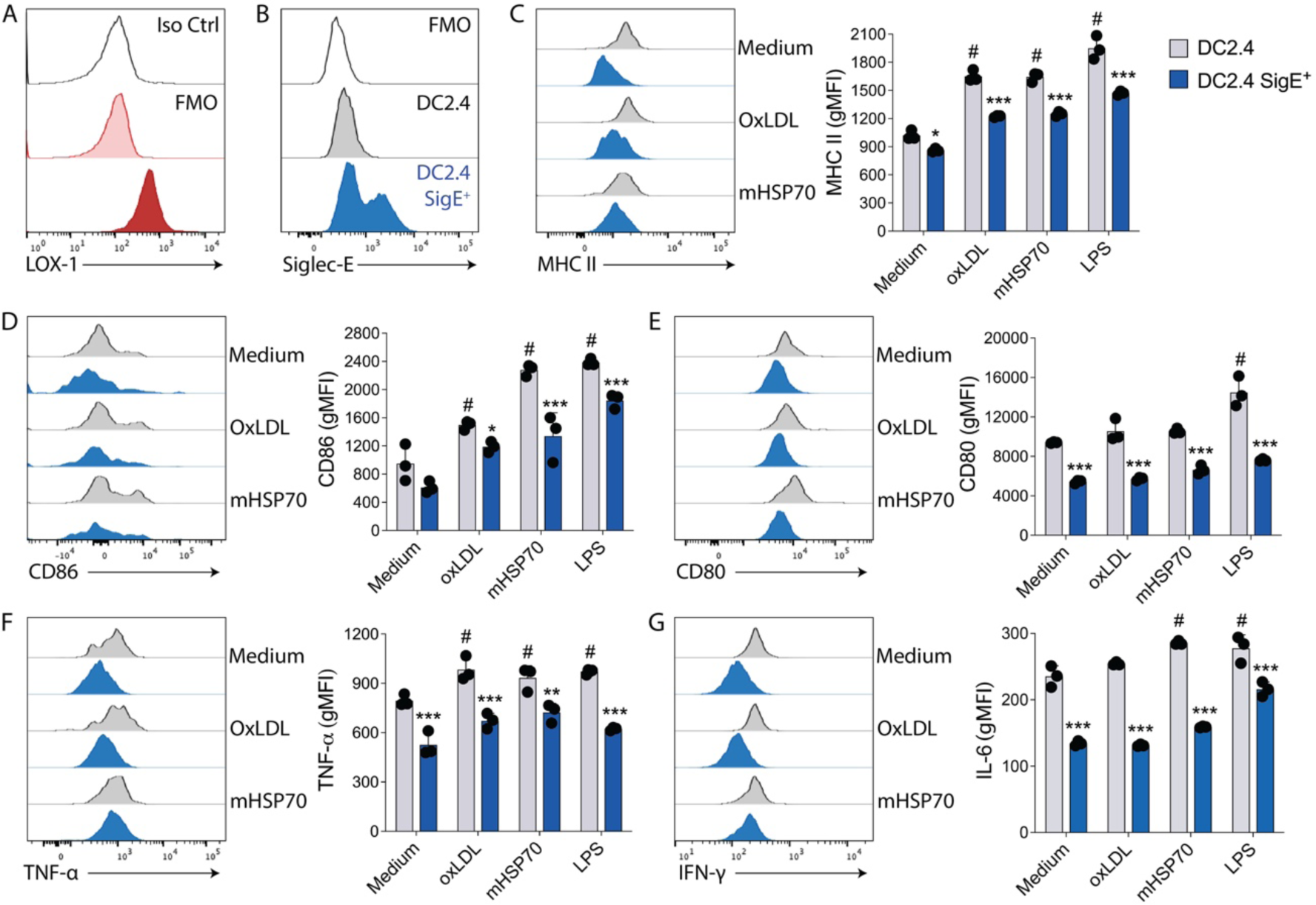
Siglec-E overexpression on DCs inhibits LOX-1-mediated activation. (**A**) LOX-1 expression in WT DC2.4 cells. (**B**) Siglec-E expression in DC2.4 cells that were lentivirally transduced with CLIP-tagged Siglec-E. WT or Siglec-E-overexpressing DC2.4 cells were treated with 50 µg/ml of mHSP70, 50 µg/ml of oxLDL or 250 ng/ml of LPS for 24h and (**C**) MHC II, (**D**) CD86, (**E**) CD80, (**F**) TNF-α, and (**G**) IL-6 levels were evaluated by flow cytometry. gMFI: geometric mean fluorescence intensity. All bars represent mean ± SD. (**C** - **G**) *p<0.05, **p<0.01, ***p<0.001 when compared to the relative WT group. Statistics by two-way ANOVA with Šídák’s post-test. #p<0.05 when compared to the WT medium group. Statistics by two-way ANOVA with Tukey post-test. Experiments are representative of three repetitions performed in triplicates.

## Discussion

Our study describes a novel innate receptor complex formed by LOX-1 and Siglec-E that mediates responses to extracellular Hsp70. Innate immunity to cell damage can induce inflammation, and even adaptive immunity ^22,23^. The magnitude of each of these inflammatory responses is highly regulated in each scenario; inflammation in response to cell damage is generally thought to lead to tissue repair rather than high-affinity antibodies or cytotoxic T memory, minimizing potential risks for autoimmunity. Tissue damage is believed to be perceived differently in the absence or presence of infection. Significant evidence supports the idea of different receptors engaged by ligands that we classify as PAMPs versus the ones we call DAMPs ^24^. This model, however, still does not indicate if receptors that evolved to recognize molecular patterns associated with cell damage can distinguish these structures from conserved microbial orthologs.

Binding of the scavenger receptor (SR) LOX-1 by human and mouse HSP70 ^9,11,25^ was previously shown to induce TNF-α and deliver antigens to antigen-presenting cells (APCs), resulting in enhancement of anti-tumor immunity and tumor regression in the absence of significant autoimmunity ^26–28^. This was rather puzzling because no regulatory mechanism had been invoked or shown that would prevent possible pathological autoimmune responses. Recently, Fong et al demonstrated that human HSP70 can bind to the human-paired receptors Siglec-5 and Siglec-14 ^8^. Their findings suggested a possible mechanism for fine-tuning of this inflammatory response: while Siglec-14 enhanced the production of IL-8 and TNF-α in human monocytes after stimulation with human HSP70, expression of Siglec-5 reduced this inflammatory signaling. Some Siglecs inhibit activatory receptors via cis interactions ^29,30^, by sequestering ligands away from their receptors and by trans-interacting with their ligands, thus regulating immune responses to self ^31^. Our results suggest that Siglec-E, an important Siglec receptor expressed in mouse innate immune cells, is regulating the downstream responses of LOX-1 when it engages mHSP70 or oxLDL. We propose that Siglec-E can control the threshold for DC activation contributing to distinguishing signal from noise triggered by self-molecules ^32^.

The responses to mHSP70 are complex and depend largely on context. For example, binding of mHSP70 to LOX-1 does not always result in T cell responses but rather consistently shows induction of TNF-α ^2,4^, and might even result in IL-10 production ^33^. This range of responses might reflect a differential expression of the Siglec-E/LOX-1 receptor complex in the relevant APCs in each experimental system. For example, TNF-α and IL-10 induction might reflect binding to LOX-1 complexed to Siglec-E, leading to inflammation and subsequent tissue repair, while binding to LOX-1 in the absence of Siglec-E association might favor the anti-tumor immunity outcomes previously reported. In light of this hypothesis, Siglec-E was recently demonstrated to interact with the scavenger receptor CD36 and control the development of atherosclerosis in mice^34^.

Finally, we have found the localization of the receptors within discrete lipid microdomains to be important for the signaling pathways triggered by the Siglec-E/LOX-1 complex. Lipid rafts are known to stabilize extensive cell surface signaling complexes, leading to the formation of immunological synapses ^35,36^. These data suggest that cell surface interaction between LOX-1 and Siglec-E may be required to stabilize both receptors in functional forms and could potentially lead to larger complexes at the cell membrane, as observed when LOX-1 binds to ox-LDL ^37^. Our data suggest that large signaling complexes, which include LOX-1, Siglec-E, and other possible receptors or associated proteins are formed during immunological responses to extracellular HSPs and perhaps other agents. Future studies should assess whether human LOX-1 has a Siglec-binding partner, such as Siglec-7 and Siglec-9, which are close human paralogs to murine Siglec-E ^38^. It was recently suggested that Siglec-E can assume a disulfide-linked dimer configuration ^39^. The homodimeric interaction is formed through a disulfide bond between cysteines at position 298, which is on the opposite side of the predicted Siglec-E/LOX-1 interface. Thus, although the interaction between Siglec-E and LOX-1 is likely between two dimers on the cell surface, the Siglec-E/LOX-1 interface would not be affected by the presence of a second Siglec-E molecule. Also, Siglec-E is itself N-glycosylated and likely carries sialic acid residues ^39^. This modification could be important for determining the interaction with LOX-1. However, the potential role of sialic acid interactions in this process has yet to be determined. Based on other examples this modification could range from being a critical ^40^ to practically unimportant ^41,42^. The composition of innate immune receptors within these complexes could explain whether a response would be biased towards inflammatory or anti-inflammatory outcomes, as well as its duration and amplitude. It also seems from our data that DC or macrophages might include both free receptors as well as Siglec-E/LOX-1 complexes^43^. Free receptors may be in transit to forming the hetero-complexes and may require ligand association to progress through membranes to form the complexes or individual receptors might have discrete stand-alone functions.

## Methods

### Cell culture

Wild-type (WT) CHO-K1 cells and all stable CHO transfectant cells were grown in Ham’s F-12 medium (Gibco) supplemented with 10% FBS (Gibco). In the case of stable transfectant CHO LOX-1 (CHO-LOX-1) clonal selection was kept at 0.4 mg/ml G418 (Gibco). Stable CHO-Siglec-E transfectant was produced through transfection with pcDNA3.1+/N-HA mouse Siglec-E plasmid. Populations of G418 (0.4 mg/ml)-resistant cells were generated after 2 weeks of cell culture. In some experiments, CHO-LOX-1 cells were transiently transfected with a Siglec-E encoding construct. All cells were maintained in a 5% CO2 humidified incubator.

WT DC2.4 cell line were grown in RPMI medium (Gibco) supplemented with 10% fetal bovine serum (Gibco), 100 U/mL of Penicillin and 100 mg/mL of Streptomycin (both from Gibco) at 37°C, 5% CO2. DC2.4 Siglec-E^+^ cell line was produced through lentiviral transduction of Siglec-E construct with CLIP tag at N-terminus. CLIP-Siglec-E was driven by the SFFV promoter.

### Animals

C57BL/6 wild-type (WT) mice were obtained from The Jackson Laboratory (Bar Harbor, ME). C57BL/6 Siglec-E KO mice described previously in ^44^ were not used because of concerns about the effect of the genomic Neo insertion. Instead, we used recently derived floxed exon-deleted Siglec-E KO mice ^45^. Both male and females were used in the experiments. All animals were between 6 to 11 weeks old.

Experiments were approved by the BIDMC Animal Care Use Committee under IACUCC0792012 and Massachusetts General Brigham (MGB) IACUC 2016N000250 and 2020N000125. All animals were housed in accordance with the Institutional Animal Care and Use Committee (IACUC) and National Institutes of Health (NIH) Animal Care guidelines.

### Purified murine HSP70 and fluorescence labeling

ADP-bound mouse Hsp70 (mHsp70) was purified from mice liver as previously described by ^10^. Purified proteins were labeled with Alexa Fluor 488 fluorescent dye using Microscale Protein Labeling Kits from Thermo Fisher, following the manufacturer’s instructions. Endotoxin levels were measured using the ToxinSensor Chromogenic LAL Endotoxin Assay Kit (Genscript), according to the manufacturer’s instructions. Only proteins from 0.2-0.45 EU/ml were used in this study. All proteins were quantitated with the Pierce BCA Protein assay kit (Thermo Scientific) before use.

### Siglec-E:Ligand ELISA interaction assay

ELISA assay was performed as an adapted protocol from ^8,46^. Briefly, wells of a 96-well plate were coated overnight at 4°C with 10 µg/ml of purified proteins in carbohydrate buffer (15 mM Na2CO3 and 35 mM NaHCO3, pH 9.6), 100µl per well. The wells were washed three times with PBS containing 0.1% Tween-20 (PBS-T) after each step. After blocking with 200 μl of blocking buffer (20 mM Tris HCl (pH7.4), 150mM NaCl, and 1% BSA) for 1h at room temperature (RT), 5 µg/ml of recombinant mouse Siglec-E-Fc chimera (R&D Systems, Cat. 5806) in 100 μl of buffer (10mM Tris-HCl (pH7.4), 150mM NaCl, 10 mM CaCl2, 1% BSA and 0.05% Tween-20) was added to each well for 2h at RT. For some experiments, 30 µg/ml of anti-LOX-1 blocking antibody (polyclonal, R&D Systems) or goat IgG isotype (Biolegend) control were added to the plate for 2h at RT before adding the recombinant Siglec-E-Fc chimera. Next, the wells were incubated with horseradish peroxidase (HRP)-conjugated goat anti-mouse IgG (polyclonal, Abcam – ab6789, 1:3000) for 1h, RT. HRP development was assayed by incubation with 3,3’,5,5’-tetramethylbenzidine (TMB) liquid substrate at RT for 30 min, and the reaction was stopped by adding 0.5N HCl per well (both from R&D Systems). Samples were analyzed at 450nm using the Benchmark Plus microplate spectrophotometer (Bio-Rad).

### Dendritic cells generation and isolation

DCs were grown from WT or Siglec-E KO bone marrows (BMDCs) in the presence of 40 ng/ml of granulocyte-macrophage colony-stimulating factor (GM-CSF) and IL-4 (both from Peprotech or Biolegend). Cells were cultured in 24-well plates in DMEM (Gibco), 10% FBS (Gibco) with pen/strip (Gibco). Non-adherent cells (DCs) were separated from adherent cells after six days in culture and were plated with 10 ng/ml of GM-CSF and IL-4. On the next day, BMDCs were incubated in media (control), or different concentrations of mHSP70 or oxLDL (Thermo Fisher) for different time points. Cells were analyzed by flow cytometry. Supernatants were collected, and cytokines were analyzed by Luminex.

CD11c^+^ cells were isolated from WT or Siglec-E KO naïve mice. Spleens were disrupted against a nylon screen and treated with Collagenase D (Roche) for 30 minutes at 37°C. Cells were labeled with anti-CD11c (clone N418)-coated magnetic beads (Miltenyi). Splenocytes were Fc blocked and CD11c^+^ cells were purified by positive selection using MACS separation columns (Miltenyi). The purity of selected cells was controlled by flow cytometry analysis. Purified DCs were analyzed by confocal microscopy.

### Binding Assays

BMDCs or splenic isolated DCs were grown on Poly-D-Lysine-coated coverslips (Corning) for 48h in serum-free AIM-V media (Gibco). Cells were incubated on ice with 10 μg/ml of Alexa 488-labeled mHSP70 or DnaK for 20 min. For receptor-mediated internalization experiments, cells were incubated with labeled mHSP70 on ice for 20 min and then at 37°C for 20 min. Coverslips were washed with PBS and fixed with 4% paraformaldehyde (PFA) for immunofluorescence experiments as described below.

Bindings analysis by flow cytometry was performed as described by ^9^. Briefly, non-trypsinized CHO cells were washed twice in PBS containing 0.5% FBS, 0.05% NaN3 and 1 mM CaCl2 (PFNC). For Siglec-E blocking, cells were incubated with 10 μg/ml of anti-Siglec-E antibody for 20 min at RT. Cells were washed twice and incubated with 10 μg/ml of labeled mHSP70 for 30 min on ice with gentle shaking. The cells were washed four times in PFNC twice and cells were analyzed in a FACS Canto II (BD Biosciences) flow cytometer.

### Immunofluorescence

After fixing with 4% paraformaldehyde, cells were permeabilized (for visualizing intracellular proteins) or not (for surface expression or binding) using 0.1% Triton X 100. Samples were blocked with 3% normal goat serum (NGS, Sigma) for 1h at RT. Cells were then stained with primary antibodies for 1h, RT and washed three times with 1x PBS. After that, cells were stained again with fluorophore-conjugated secondary antibodies for 1h at RT. After three washes with 1x PBS, cells were stained with 1µg/ml of Hoechst 33342 (Thermo Scientific) in PBS for 10 min at RT. Fluorophores were visualized using the following filter sets: 488 nm excitation and bandpass 505–530 nm emission filter for Alexa 488; 543 nm excitation and bandpass 560–615 nm for Cy3/Alexa 594 nm; and 633nm excitation, in Zeiss Confocal Microscope. Images were processed using ZEN 2 blue edition and Abode Photoshop. Primary antibodies used were: goat anti-mouse Siglec-E (polyclonal IgG, R&D Systems) and mouse anti-cell membrane Hsp70 (clone 1H11, StressMarq) were used as 1:100, rat anti-LOX-1 (polyclonal, from Dr. Sawamura, described in ^47^ was used as 1:200, mouse anti-HA antibody (clone 16B12, Covance) as 1:250, rabbit anti-caveolin (polyclonal, Sigma-Aldrich) and mouse anti-c-Myc (clone 9E10, Biolegend) were used as 1:300; rabbit anti-HA (cat H6908, Sigma) was used as 20 µg/ml. All secondary antibodies were purchased from Jackson ImmunoResearch and used as 1:300. All antibodies were prepared in 3% NGS solution.

### Deglycosylation of mHSP70

mHSP70 or fetuin (New England Biolabs, positive control) were subjected to cleavage by PNGase F (New England Biolabs), following the manufacturer’s instructions. Briefly, for each reaction, 20 μg of mHSP70 and fetuin were denatured at 100°C for 10 min and then mixed with Glycobuffer, 10% NP-40, and 1 μl of PGNase F, during 1h at 37 °C. For protein detection, the gel was stained using Coomassie Blue (Bio-Rad).

### Immunoprecipitation

WT splenocytes were incubated with 10 µg/ml of mHSP70, or media for 30 min at 37°C. For some experiments, spleen cells were pre-treated with 20 mM of the cholesterol-sequestering agent methyl-beta-cyclodextrin (MβCD, Sigma) for 4h. After that, Siglec-E was immunoprecipitated. Spleen cells lysates were prepared in NP-40 lysis buffer (containing 1% Nonidet P-40, 150 mM NaCl, 1mM EDTA, 1 mM PMSF - Boston BioProducts) with 1x Halt protease inhibitor cocktail (Thermo Scientific). Samples were pre-cleared with 40 µl of protein G Sepharose beads (50% slurry, GE Healthcare) plus 5 μg Goat IgG (Invitrogen), overnight at 4°C with rotation. Protein concentration was measured with BCA assay and 350 µg of lysates were incubated with 5 μg of anti-Siglec-E antibody (polyclonal goat IgG - R&D Systems – cat. AF5806) or goat IgG for 2h at 4°C, and then 40 μl of 50% bead slurry was added for overnight at 4°C with rotation. The beads were washed four times with lysis buffer and complexes were eluted by boiling in Laemmli sample buffer (Bio-Rad) for western blot.

### Western Blot

For Western blotting, 30 μg of protein were resolved by 4–15% gradient SDS-PAGE (Bio-Rad) and transferred to PVDF (polyvinylidenefluoride) membranes. Membranes were blocked with 5% non-fat milk and immunoblotted with primary antibodies. After washing, membranes were incubated with secondary antibodies that are HRP-conjugated. The membrane reactions were visualized by Perkin Elmer enhanced chemiluminescence reagents. Primary antibodies: goat anti-mouse Siglec-E (polyclonal IgG, R&D Systems) and mouse IgM anti-mouse Siglec-E (clone F-7, Santa Cruz) were used as 1:200, rabbit anti-LOX-1 (clone EPR4025, Abcam), and mouse anti-HSP70 (clone C92F3A-5, StressMarq) were used as 1:10000. Secondary HRP-conjugated antibodies: goat anti-mouse IgM (Santa Cruz, cat. Sc-2064) was used as 1:2000, horse anti-mouse IgG (cat. 7076, Cell Signaling) was used as 1:3000, mouse anti-rabbit IgG, light chain-specific (clone 5A6-1D10, Jackson ImmunoResearch), and goat anti-rabbit IgG (cat. 7074, Cell Signaling) was used as 1:3000.

For western blotting with the lectin, the membranes underwent a 30 min-blocking at RT using 1x carbo-free blocking solution (Vector) containing 1% Tween-20 (Sigma). The membranes were then blotted with 1 μg/mL biotinylated Sambucus Nigra Lectin (SNA, EBL, Vector) in in the TBS with 0.1M Ca2^+^ and 0.1M Mg2^+^ for 1h at RT. Following three washes with PBS 1x with 0.05% Tween-20, membranes were incubated with goat anti-biotin-HRP at 1:5000 (Vector) in PBS 1x. Sialyated proteins on the membranes were visualized by PierceTM ECL Western Blotting substrate solution (ThermoFisher).

### Flow cytometry

Cells were initially washed with PBS and stained with Fixable Viability Dye eFluor 780 (eBioscience) by the detection of dead cells. Cells were then Fc blocked for 20 minutes on ice and surface markers were stained by incubation for 30 minutes with antibodies in PBS with 2% fetal bovine serum (FBS). Intracellular staining was performed after treating cells with the eBioscience® Foxp3 Fixation/Permeabilization solution, followed by incubation with antibodies for 1h at RT. The following antibodies were used for murine cells: I-A^b^ (MHC II, clone AF6-120.1; 1:200), CD86 (clone GL1; 1:100), CD11c (clone HL3; 1:100), CD45R/B220 (clone RA3-6B2; 1:200), CD11b (clone M1/70; 1:100) from BD Biosciences; LOX-1 (clone 214012, 1:50) from R&D Systems; Siglec-E (clone M1304A01, 1:200), IL-6 (clone MP5-20F3, 1:100), TNF-α (clone MP6-XT22, 1:66) and CD80 (clone 16-10A1, 1:100) from Biolegend.

For intracellular signaling pathway analysis, cells were fixed with 4% PFA for 10 min at RT followed by permeabilization with the True-Phos Perm buffer (Biolegend) for 20 min on ice, followed by incubation with antibodies for 30 min at RT. The following antibodies were used for the signaling pathways: phospho-Src (Tyr418, clone SC1T2M3; 1:10) from Thermo Fisher, mouse IgG2b isotype control from Biolegend, SHP-1 (clone E1U6R, 1:50) and rabbit IgG isotype control from Cell Signaling. Cells were analyzed using FACS Canto II or Fortessa X-20 (both from BD Biosciences). Data obtained were analyzed using Flowjo software (version X, Tree Star).

### Cytokines measurements

Cytokine profiles were determined from culture supernatants using a ProcartaPlex 11-Plex Kit from Thermo Fisher Scientific, containing the following cytokines: IL-6, TNF-α, IL-12p70, IL-10, IL-1β, IL-10, IL-18, GM-CSF, IL-4, IL-5, IL-13, and IFN-γ. Data are expressed as Log_10_ concentration (pg/ml) in a heatmap created with the Morpheus matrix visualization and analysis tool from the Broad Institute (https://software.broadinstitute.org/morpheus).

### Protein Homology modeling and Molecular docking

The proteins LOX-1 (Uniprot ID: Q9EQ09) and Siglec-E (Uniprot ID: Q91Y57) were modeled using Modeller v9.10 software ^48^. Both proteins were initially submitted to the same homology modeling protocol. Briefly, we first searched for three-dimensional templates using BLAST, HHPRED and PHYRE2 servers ^49,50^. Then we performed a structural alignment using selected templates against LOX-1 or Siglec-E sequences through TCOFFEE server (Notredame, C. *et al*, 2000) in order to choose the best template(s). After that a basic Modeller algorithm generated 100 models. The best model was chosen according to DOPE score, Ramachandran plot analysis, QMEAN score ^52^ and ModFOLD6 score ^53,54^ of each residue. When necessary, a loop refinement was used to correct loop regions with bad modeling scores. The protein-protein interaction was assessed through ClusPro 2.0 ^15^. The parameters were maintained as default. The electrostatic potential of the structures was computed with DelPhi software (available in http://honig.c2b2.columbia.edu/) and analyzed with UCSF Chimera and UCSF Chimera X softwares ^55,56^.

### Statistics

For the comparison of two independent groups, we used an unpaired Student’s t-test for comparison. For the multiple-group comparison, one-way or two-way analysis of variance (ANOVA) was used to determine differences. We used Tukey post-hoc tests for multiple comparisons between levels for one-way ANOVA, and Sidak’s multiple comparisons test for two-way ANOVA. Significant differences were set for p ≤ 0.05. After Kaplan-Meier survival curves were generated, a log-rank test was used for statistical inference between experimental groups. Prism software was used for statistical analysis and plotting graphs (GraphPad Software, Inc.).

## Data availability

Further information and requests for resources and reagents should be directed to and will be fulfilled by the lead contact.

## Acknowledgments

We want to thank Dr. Ajit Varki from the UC San Diego for useful materials and critical discussions, Dr. Jawahar Mehta from the University of Arkansas for Medical Sciences for provided mice, Dr. Tatsuya Sawamura from the Shinshu University for provided antibody, Lay-Hong Ang from the BIDMC Confocal Imaging Core for the assistance in immunofluorescence experiments, and Sylvain Lehoux from the BIDMC Glycomics Core for the assistance in glycosylation experiments.

## Contributions

T.J.B., A.M., Y.Z., M.M.R., E.H., L.V.R., C.B., and S.K.C. designed the research. T.J.B., K.L. and A.M., performed binding assays. T.J.B. and A.M. performed immunofluorescence. T.J.B. and S.S.S. performed ELISA assays. T.J.B., K.L., and A.M.. T.J.B., K.L., and I.T.L. performed dendritic cell cultures. T.J.B. and K.L. performed flow cytometry and Luminex experiments. Y.Z. created DC2.4 cell lines. T.J.B., A.M., and B.J.L. created CHO cell lines. I.T.L. maintained all the cell lines. M.M.R. performed in silico docking analyses. T.J.B., L.V.R., C.B., and S.K.C. wrote the manuscript. All authors edited the manuscript.

