## Supplementary Figures and Tables for "Innate extracellular Hsp70 inflammatory properties are mediated by the interaction of Siglec-E and LOX-1 receptors"

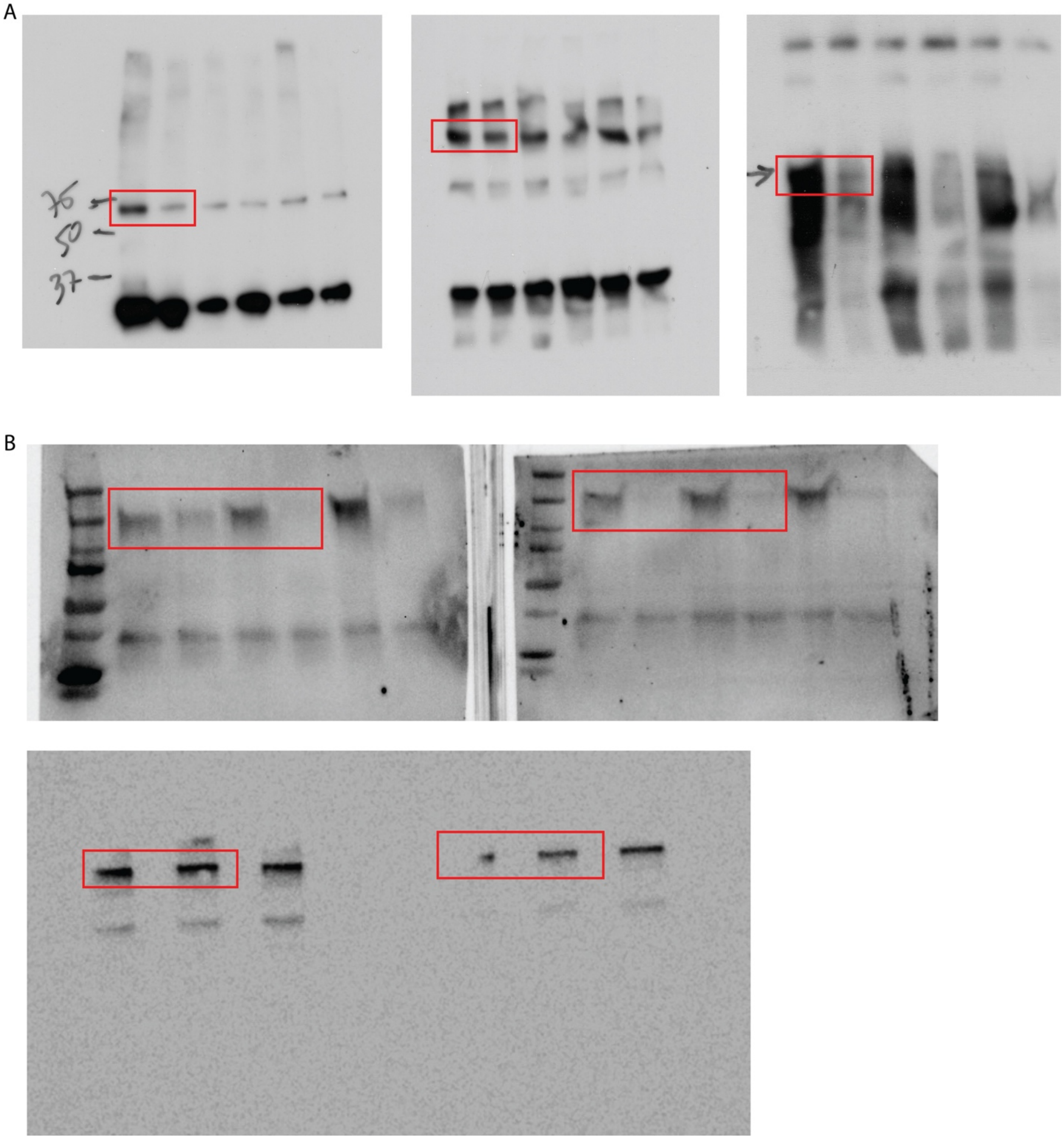


**Supplementary Fig. 1. Uncut Western blot membranes.** (**A**) Membranes from Fig 3D. (**B**) Membranes from Fig 5F.

### Supplementary Tables

**Supplementary Table 1. Residues interact through hydrogen bonds between a Siglec-E molecule and a LOX-1 dimer.**

| Siglec-E | LOX-1 |
|---|---|
| LYS32 | GLN111<br>ASP112 |
| ARG34<br>THR36 | GLU118 |
| ASP37 | GLN111<br>TYR109 |
| THR39 | ILE123 |
| ARG75 | GLU255 |
| ASN76 | TYR239 |
| TRP85 | ASP112 |
| ARG86 | ASP112<br>SER60 |
| TRP159 | SER240<br>ASN256 |

**Supplemental Table 2:** Comparison of colocalization coefficients between Siglec-E and LOX-1 on untreated or mHSP70-treated murine dendritic cells[a]

| **Coefficients** | **Siglec-E-LOX-1 colocalization** | | |
| --- | --- | --- | --- |
| | **Untreated** | **mHSP70** | ***p* value** |
| Pearson's correlation | 0.347 ± 0.078 | 0.543 ± 0.0.45 | *0.0157*[b] |
| Spearman's rank correlation value | 0.383 ± 0.033 | 0.685 ± 0.041 | *0.0005*[b] |

[a] Results are means ± standard deviation of coefficients from five analyzed areas
[b] Untreated vs mHSP70
by two-way ANOVA with Sidak's multiple comparisons test
